# A CXCR3-activating peptide increases Tear Break Up Time and corrects corneal haze in a rabbit model of environmental dry eye

**DOI:** 10.1101/2022.12.07.519513

**Authors:** Alan Wells, Yadong Wang, Hanshuang Shao, Peri Sohnen, Shivalingappa K. Swamynathan

## Abstract

**Purpose:** Environmentally-triggered dry eye disease (DED) or keratoconjunctivitis sicca (KCS), which constitutes the majority of DED cases, currently is palliatively treated with aqueous replacement solutions that do not target the dysfunction of the mucin and lipid components of tears. Herein, we tested whether a peptide that increased goblet cell numbers in a model of scleral chemical injury would also improve tear quality in environmental DED.

**Methods:** Environmental DED was established by exposing New Zealand white rabbits (8 per group, female) to 20% humidity with rapid air replacement and b.i.d. atropine sulfate eyedrops for 3 weeks prior to test article administration; this continued for the subsequent 3 weeks of testing. Animals were dosed by (A) saline, (B) b.i.d. eyedrop of peptide in saline, (C) b.i.d. eyedrop of peptide in coacervate, or (D) weekly subconjunctival injection of peptide.

**Results:** The environmental DED was established with both Schirmer and TBUT being reduced by from baseline at the start of test article; these levels were maintained as low through the testing period. All three treatment regimens increased TBUT approximately 3x to levels greater than prior to desiccation, with little effect on Schirmer. Corneal haze was present in all eyes after induction, and largely cleared up by all three treatments. End of study enucleation of the eye did not show any changes in goblet cells numbers, which remained high throughout the induction and treatment.

**Conclusions:** The treatment of environmental DED/KCS with a peptide that activates CXCR3 improved tear quality and reversed corneal pathology by promoting tear stability, while not affecting aqueous volume of the tears.

## Introduction

Dry eye disease (DED) or technically known as keratoconjunctivitis sicca (KCS) is a syndrome with a number of causes [1–3]. The basal phenotype of insufficient tear coverage of the sclera is often treated palliatively with physical replacement of tears. The only treatments that target a cause of DED are those that ameliorate auto-immune disease related inflammation [4], however these immunosuppressive agents are germane against only about 1 in 10 cases of DED. This leaves the vast majority of cases with only temporary symptomatic relief.

Regardless of the mechanistic cause, the tissue manifestations of DED converge on damage to the sclera of the eye [2, 5]. Once this is compromised, the goblet cells malfunction and are eventually lost due to the non-specific inflammatory damage. This leads to a decrease in the mucin layer of the tears, with resultant inability of the aqueous layer to adhere to the scleral surface [1]. Once the goblet cell mucin is insufficient, additional aqueous components are temporary, leading to either frequent administration or even the paradoxical watery/teary dry eye symptoms.

Serendipitously, we found that activators of the CXCR3 receptor can provide for goblet cell rescue in a rabbit model of toxic dry eye [6]. After trabeculectomy, subconjunctival injection of a peptide activator of CXCR3 led to an increase in number of goblet cells at 21 days to a level higher than prior to surgery. Even in the face of mitomycin C droplets, the goblet cells rebounded to match pre-surgical levels. This is intriguing due to the recent focus on the centrality of mucins in DED [5, 7]. Based on these findings, we investigated whether CXCR3 activators could alleviate dry eye metrics in causes of DED in addition to surgical and toxic injury. Herein, we report that a peptide activator of CXCR3 was able to restore mucin/lipid functioning in a rabbit model of environmental DED, as measured by prolongation of Tear BreakUp Time (TBUT).

## Methods

### Materials

The peptide used was a 22-mer (OH-PESKAIKNLLKAVHKEMSKRSP-NH2) manufactured by PolyPeptide Inc (San Diego, CA, USA) to >95% purity as a lyophilized powder. This was delivered in three different formulations. For subconjunctival injection, 3ug were dissolved in 100ul of sterile PBS for weekly injections. For eyedrops, 10ug were dissolved in 50ul of sterile PBS for twice daily drops (BID). For the slow release formulation, 10ug were dissolved in 50ul of coacervate for BID inoculation. The control was 50ul of sterile PBS applied BID.

The coacervate is a Heparin and PEAD (1:3.6) mixture with each component at a 10mg/mL concentration in water [8, 9]. The peptide was added at a 1:10 dilution in PBS to the final concentration.

### Study Design

The environmental dry eye model was tested under contract at Absorptions Systems (San Diego, CA, USA). This CRO follows the ARVO guidelines for animal research, and the study was approved by the ASC IACUC prior to initiation of the study. ASC is AAALAC-accredited site, and is in compliance with ASC’s Animal Welfare Assurance (D16-00645 [A4282-01]) filed with the NIH.

Female New Zealand White Rabbits (8 per group) were used at 125-175 days old (3-5 kg) at the time of induction. For induction, all animals were dosed with Atropine Sulfate (1%) in both eyes (OU) 2x daily (BID) for 2–3 weeks prior to test article administration to induce KCS. In addition, humidity in the animals’ housing room was maintained at ~20% throughout the study to promote eye dryness. Non-responder animals were removed from the study at the end of the induction phase. This procedure was continued for all animals remaining on study until study completion.

The study was conducted as outlined in Table 1 (attached draft report). This was done in two staggers with half the rabbits in each group (4) being treated in each stagger.

**Table 1.**
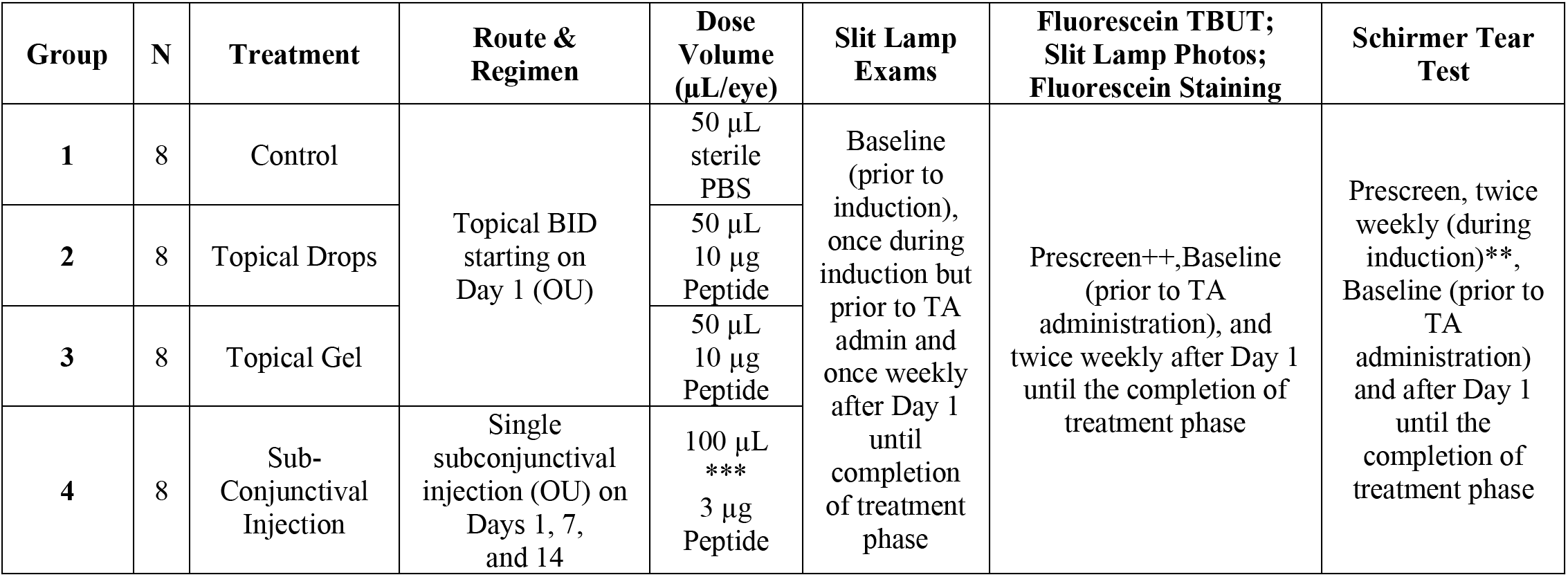
Study Design Table.

## Results

The main measurements relevant to tear function were the Schirmer Tear Test (STT) and TBUT times. For the STT, the baseline values of 12-15mm were reduced minimally to 9-12mm after induction, and tended to drift lower during treatment to 8-11mm, regardless of the delivery mode (Table 2). Thus, the amount of aqueous tear was uncorrected by the CXCR3 activating peptide.

**Table 2.** STT values prior to induction, prior to treatment and after treatment.

| Treatment | Day -21 |  | Day 0 |  | Day 21 |  | P Value |
| --- | --- | --- | --- | --- | --- | --- | --- |
|  | OD | OS | OD | OS | OD | OD |  |
| Saline Ctrl | 15.5±3.2 | 13.1±2.9 | 12.1±3.6 | 9.0±2.9 | 8.4±2.2 | 8.3±3.0 | n.s. |
| Eyedrop | 14.6±5.6 | 14.0±2.8 | 11.3±3.3 | 9.9±3.7 | 9.9±3.9 | 7.9±2.0 | n.s. |
| SR Peptide | 14.6±3.3 | 13.0±4.1 | 9.8±3.4 | 9.3±3.2 | 9.4±2.9 | 8.4±1.8 | n.s. |
| ConjInj | 12.6±2.8 | 12.5±2.7 | 10.3±2.5 | 8.8±2.7 | 11.0±1.9 | 9.8±2.0 | n.s. |

The TBUT was altered much more by the induction and treatments. Initial TBUT times of 8-9s were reduced to half (3.5-5s) after induction and prior to treatment (Table 2; supplemental data). TBUT remained low in the saline treated eyes, but rebounded to baseline in the first week of treatment of all three deliveries (Figure 1). Interestingly in all three treatment groups, the TBUT exceeded baseline by the end of week 3 (Table 3). This increase in TBUT was also noted when analyzing each stagger separately.

**Figure. 1.**
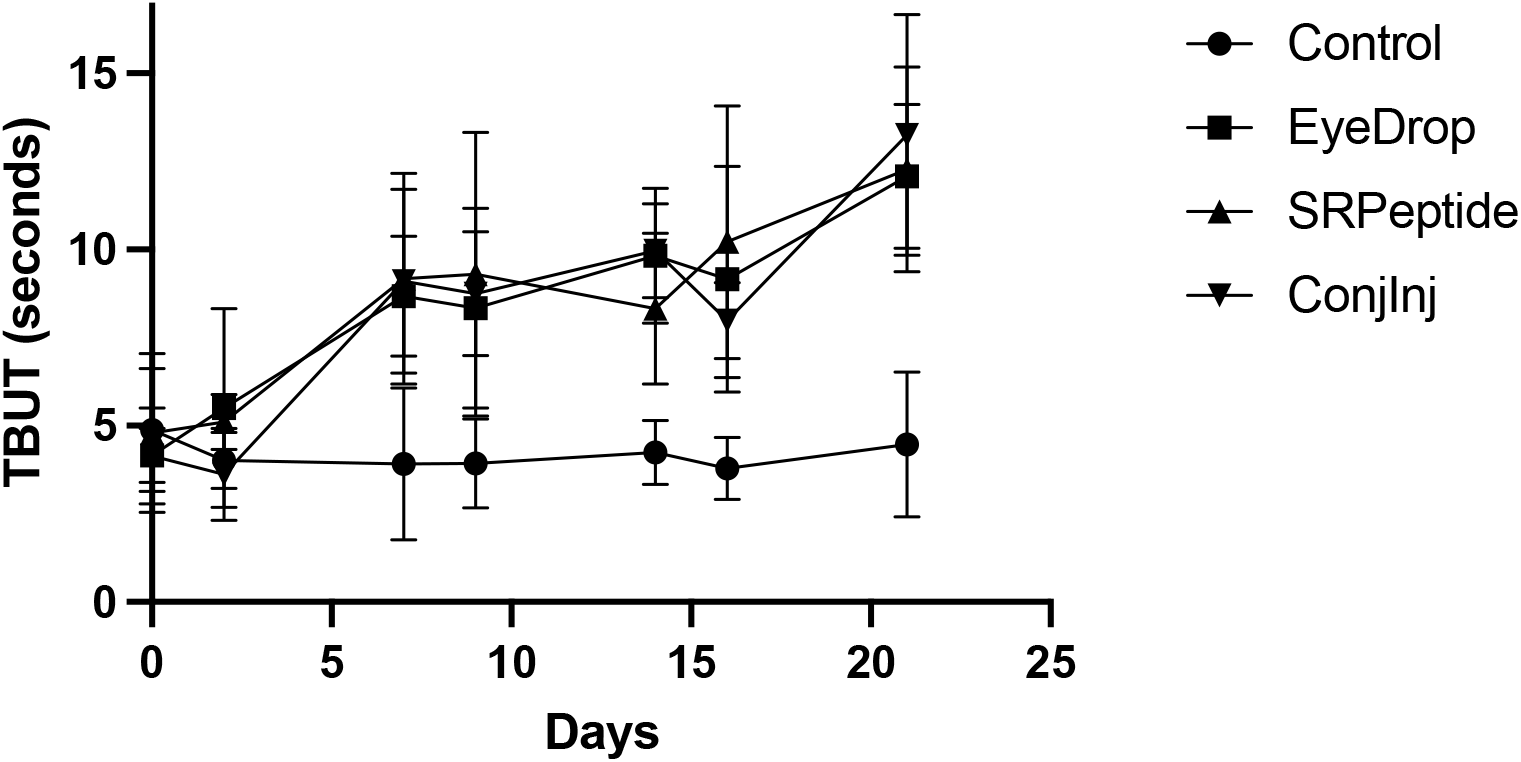
Increase in TBUT upon treatment of rabbits. Rabbits (8 per group) were treated with the peptide weekly in subconjunctival injection formulation, or twice per day in saline eyedrops or in a coacervate formulation. The control was saline eyedrops twice per day. On days 2, 7, 9, 14, 16, and 21 the eyes were assessed for TBUT and Schirmer. Presented are averages, with all treatments groups being statistically greater than saline control by day 7 (p < 0.05 using t-Test comparisons).

**Table 3.** TBUT values prior to induction, prior to treatment and after treatment.

| Treatment | Day -21 |  | Day 0 |  | Day 21 |  | P Value |
| --- | --- | --- | --- | --- | --- | --- | --- |
|  | OD | OS | OD | OS | OD | OD |  |
| Saline Ctrl | 8.6±2.7 | 8.3±1.7 | 4.9±2.2 | 4.7±1.2 | 4.8±1.7 | 4.1±2.1 |  |
| Eyedrop | 8.7±2.0 | 8.4±1.4 | 4.0±0.6 | 4.7±1.3 | 11.3±4.2 | 10.9±4.7 | >0.01 |
| SR Peptide | 7.9±2.0 | 7.9±1.7 | 4.7±1.8 | 4.7±1.6 | 13.3±4.1 | 12.7±5.1 | >0.01 |
| ConjInj | 9.0±3.9 | 8.5±2.3 | 3.7±0.7 | 4.9±2.3 | 13.2±5.8 | 12.7±5.3 | >0.01 |

The increase in the TBUT combined with no effect on STT suggested that either the mucin or lipid layer was altered, in accordance with the earlier findings on an increase in goblet cells after toxic (mitomycin-C) injury [6]. Thus, we asked whether the peptide altered the goblet cells. Histopathological examination of the enucleated globes at the end of the study (day 21 of treatment) found that goblet cell density was similarly high in all treatment and control groups (data not shown).

However, the increase in mucin production did appear to alleviate the desiccation-induced corneal pathology. In all groups, all 16 eyes displayed corneal pathology at the end of the three-week induction period. At the end of the three-week test article administration the control group still had corneal pathology in all 16 eyes. However, the number of affected eyes were reduced after treatment with eyedrops (4 of 16 eyes showing pathology), slow-release gel formulation (2 of 16) and subconjunctival injection (6 of 16). All treatments were statistically different from the saline-treated control (at P < 0.05).

## Discussion

Dry eye disease results from a plethora of insults to the surface of the eye [1–3], with only a small number of auto-immune conditions having disease modifying treatments (mainly nonspecific immune suppression) [4, 10]. The vast majority of the cases are treated symptomatically. When the insult is not a singular episode such as a toxic substance or ocular surgery [11], this leads to a chronic condition. As environmental causes, relative low humidity, are impractical to avoid for most patients, new treatments that have the potential to modify the course of the disease are needed.

The pathophysiologies of DED eventually converge on damage to the mucin-producing goblet cells [1, 7] that in turn leads to the inability of tears to adhere to the ocular surface. This unifying principle in DED impels us to seek treatments that improve goblet cell functioning and survival. To that end, we serendipitously found that a peptide activator of CXCR3 increased the number of goblet cells after ocular surface insult during trabeculectomy [6]. Herein, we asked whether this same stimulus could reverse dry eye due to environmental insult. That the TBUT was restored and even bolstered above baseline without an improvement in STT suggested that the peptide could address this type of DED. While we could not enumerate an increase in goblet cells likely due to the high numbers present even after 6 weeks of desiccation treatment.

The effects on goblet cell number and function need to be pursued in ancillary studies. This could take the form of laboratory investigations into goblet cell progenitor (stem cell) survival, proliferation, and functioning. While these studies are underway, the fall outside the purview of this initial report, as the translation of in vitro effects to animal interventions is far from straightforward.

There are a number of caveats in translating these findings to the human condition, not the least being that rabbit eyes have high densities of goblet cells and robust TBUT. Thus, whether such enhanced functioning will transfer to the human condition or to similar extent will require trials in patients. Second, loss of the aqueous component of tears may remain a barrier in many patients for whom tear volume is a challenge. In such cases a combinatorial approach would make intuitive sense. For situations in which ongoing auto-immune or allergic inflammation initiates the ocular surface damage, just supporting the goblet cells is unlikely to be disease modifying and also require combinatorial therapies. Despite these concerns, it is only upon testing in patients can we address whether this approach represents a new quiver in our armamentarium against dry eye disease.

## Acknowledgements

This study was supported by funds from the University of Pittsburgh PreMIC Competition, supported by the Richard King Mellon Foundation. Neither the PreMIC Competition nor the Foundation had input into the design, interpretation or communication of this experiment and manuscript.

## Conflict of Interest Statement

AW, YW and SKS have filed a provisional patent related to this treatment, with assignment to their respective institutions. AW has an IP position on the peptide and YW has an IP position on the coacervate. AW is a founder, CSO and shareholder in Ocugenix Inc, a preclinical start biotech company that has an interest in licensing this intellectual property.

